# Outer membrane vesicles produced by pathogenic strains of *Escherichia coli* block autophagic flux and exacerbate inflammasome activation

**DOI:** 10.1101/2021.04.20.440604

**Authors:** Laure David, Frédéric Taieb, Marie Pénary, Pierre-Jean Bordignon, Rémi Planès, Salimata Bagayoko, Valérie Duplan-Eche, Etienne Meunier, Eric Oswald

## Abstract

*Escherichia coli* strains are responsible for a majority of human extra-intestinal infections, resulting in huge direct medical and social costs. We had previously shown that HlyF encoded by a large virulence plasmid harbored by pathogenic *E. coli* is not a hemolysin but a cytoplasmic enzyme leading to the overproduction of outer membrane vesicles (OMVs). Here, we show that these specific OMVs inhibit the autophagic flux by impairing the autophagosome – lysosome fusion, thus preventing the formation of acidic autophagolysosome and autophagosome clearance. Furthermore, HlyF-associated OMVs are more prone to activate the non-canonical inflammasome pathway. Since autophagy and inflammation are crucial in the host’s response to infection especially during sepsis, our findings reveal an unsuspected role of OMVs in the crosstalk between bacteria and their host, highlighting the fact that these extracellular vesicles have exacerbated pathogenic properties.

## Introduction

Although the majority of *E. coli* strains are commensal bacteria colonizing the digestive tract, some are pathogenic because they have acquired an arsenal of virulence factors that allow them to overcome innate or acquired defense mechanisms of their host (Sarowska et al., 2019). HlyF is a bacterial protein encoded by a gene present on a virulence plasmid found in strains of *E. coli* responsible for extraintestinal infection in both humans and animals (Peigne et al., 2009). More specifically, it is found in strains responsible for neonatal meningitis, avian colibacillosis (Johnson et al., 2008; Kim et al., 2020; de Oliveira et al., 2015) and more recently in an emerging serovar of enterohemorrhagic *E. coli* (EHEC) responsible for hemolytic and uremic syndrome in human (Soysal et al., 2016). The *hlyF* gene does not code for hemolysin as its name would suggest, but for a bacterial cytoplasmic protein that contributes to the bacterial virulence (Murase et al., 2016). The HlyF protein displays a predicted catalytic domain of the short-chain dehydrogenase/reductase superfamily. This conserved domain is involved the ability of HlyF to promote the production of bacterial OMVs (Murase et al., 2016). Noteworthy, HlyF protein is not detected in OMVs. OMVs are nanoparticles consisting of a lipid bilayer envelope including membranous proteins and LPS originating from the parental strain and containing luminal cargos (Jan, 2017; Schwechheimer and Kuehn, 2015). Similarly to eukaryotic exosomes and microvesicles, OMVs are a means of communication between bacteria and eukaryotic host cells (Kuehn and Kesty, 2005; Kulp and Kuehn, 2010). Interestingly, we have observed that eukaryotic cell treatment with culture supernatants of *E. coli* strains expressing HlyF induced LC3-positive vesicles accumulation in host cells. At that time, we thought we were observing an exacerbation of autophagy and we interpreted this phenotype as related to the greater quantity of OMVs present in the supernatant (Murase et al., 2016).

Here, we show that extracellular vesicles from *E. coli* strains producing HlyF have exacerbated pathogenic properties compared to OMVs produced by isogenic strains unable to produce a functional HlyF. Our results revealed that these OMVs inhibit the autophagic flux and thus are more prone to promote the activation of the non-canonical inflammasome pathway, the main host defense mechanism against infections that must be tightly regulated to ensure a correct antimicrobial response (Biasizzo and Kopitar-Jerala, 2020; Feng et al., 2019; Ho et al., 2016; Qiu et al., 2019). Thus, these OMVs, produced by bacteria responsible of extraintestinal infections, could destabilize the host response to the infection, thus facilitating bacterial pathogenesis.

## Results

### *OMVs from HlyF-expressing* E. coli *specifically induce an increase of LC3-positive vesicles numbers*

In agreement with our previous results (Murase et al., 2016), bacterial expression of wild type (WT) HlyF is associated with an overproduction of OMVs (**Fig. 1A**). Moreover, HeLa cells treatment with culture supernatant of these HlyF WT strains induces an increase of LC3-positive structures number in host cells, as denoted by vesicles-associated LC3 foci (**Fig. 1B**). To take into account a dose effect due to overexpression of OMVs in bacteria producing HlyF, eukaryotic cells were then treated with the same number of purified OMVs from isogenic *E. coli* strains expressing or not HlyF. To do so, we analyzed OMVs by transmission electron microscopy (TEM) and checked that the concentration of OMVs is correlated to protein concentration (data not shown). At the same concentration, only OMVs from the wild type strain expressing HlyF specifically induce an increase of LC3-positive structures number in host cells (**Fig. 1C**). We repeated these experiments using the same amount of OMVs purified from the laboratory *E. coli* strain BL21(DE3) devoid of virulence factors and flagellin, expressing wild-type or inactivated HlyF (**Suppl. Fig 1A**). OMVs were analyzed by TEM (**Fig. 1D**), but also by dynamic light scattering (**Suppl. Fig. 1B**) and we performed protein and LPS dosages (**Suppl. Fig. 1C**) to refine their characterization. These results showed that OMVs produced from both strains expressing active and inactive HlyF have a similar diameter (25.95nm +/- 4.91 and 24.5nm +/- 4.86 respectively) and comparable concentration of protein and LPS per vesicle. The increase of LC3-GFP-associated LC3-positive foci (**Fig. 1E**) as well as the significant increase of LC3-II level (**Fig. 1F-G**) confirmed the specific HlyF-dependent increase of LC3-positive structures number following cell treatment with OMVs from BL21-producing wild type HlyF.

**Figure 1.**
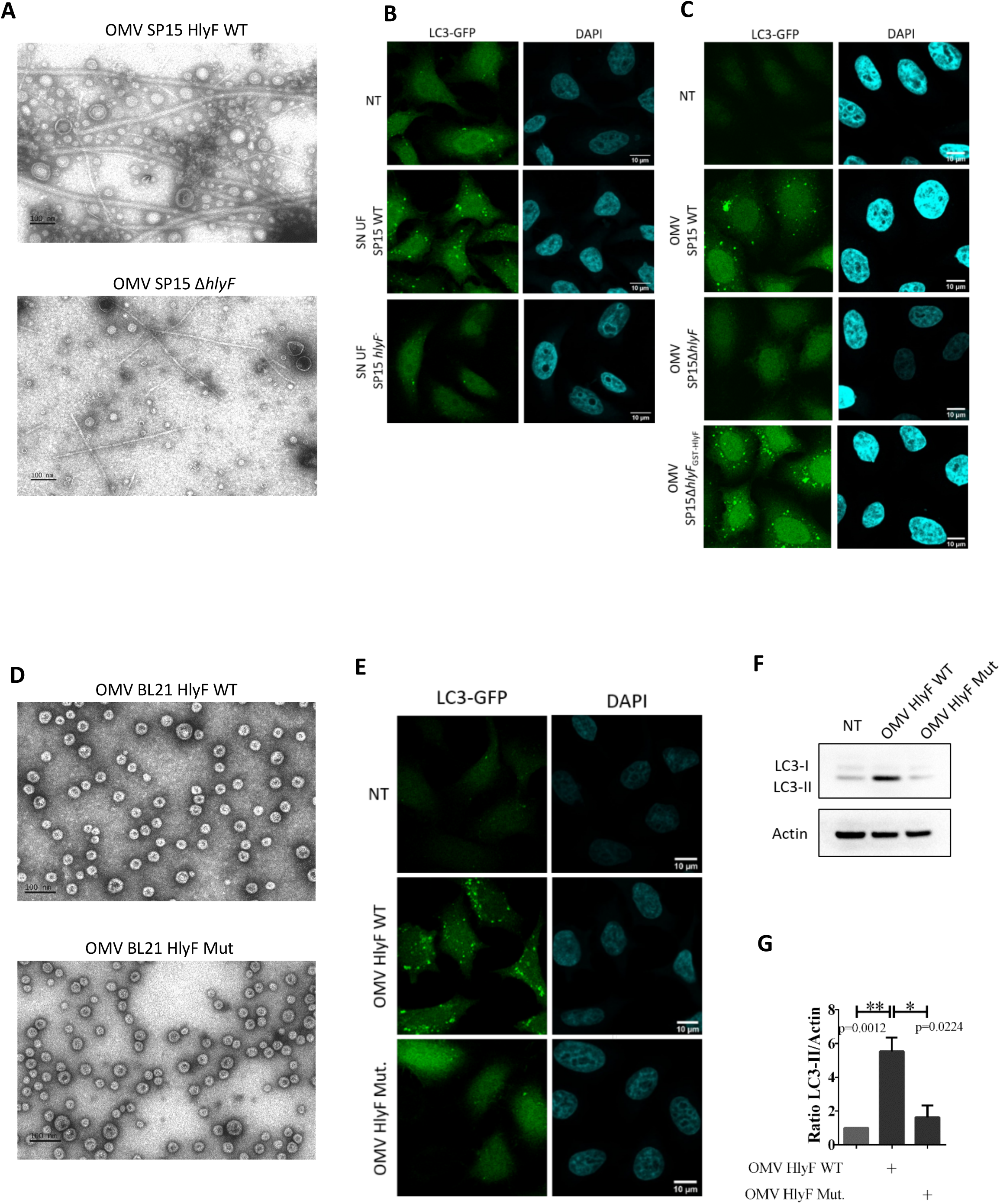
Increased LC3-positive structures numbers in epithelial cells treated by OMVs from *E. coli* producing HlyF. (**A**) Images of purified OMVs with equivalent volumes from wild type SP15 (SP15 WT) and from SP15Δ*hlyF* visualized by negative staining transmission electron microscopy (TEM). Scale bar = 100nm. Images representative of 3 independent experiments. (**B-C**) Images of confocal microscopy of LC3-GFP (green) and DAPI (blue) in LC3-GFP-Hela cells treated 3hr with supernatant from wild type SP15 (SP15 WT) and SP15Δ*hlyF* 250μL/mL after ultrafiltration (**B**) or with purified OMV with equivalent protein dosage from wild type SP15 (SP15 WT) 10μg/mL, SP15Δ*hlyF* 10μg/mL and SP15Δ*hlyF*_GST-HlyF_ 1μg/mL (**C**). Scale bar = 10μm. Images representative of 3 independent experiments. (**D**) Images of purified OMVs with equivalent protein dosage from BL21 expressing HlyF WT or inactivated (Mut) visualized by negative staining TEM. Scale bar = 100nm. Images representative of 3 independent experiments. (**E**) Images of confocal microscopy of LC3-GFP (green) and DAPI (blue) in LC3-GFP-Hela cells treated 3hr with OMV BL21 HlyF WT or OMV BL21 HlyF Mut 5μg/mL. Scale bar = 10μm. Images representative of 3 independent experiments. (**F**) Western blot analysis of LC3 and actin in cell extracts of Hela cells treated 3hr with OMV from BL21 HlyF WT and OMV BL21 HlyF Mut 5μg/mL. Experiment reproduced 3 times independently. (**G)** Quantification of the LC3B-II/actin ratios of the Fig. 1F obtained by densitometric analysis of 3 independent experiments. The graph shows the mean and the standard deviation for each condition. ** p<0,01 and *p<0,05 t test.

### Increased LC3-positive vesicles numbers induced by OMVs from HlyF-expressing E. coli is dose-dependent

To quantify the number of LC3-positive foci per cell, we counted the fluorescently labelled GFP-LC3 puncta per cell using the imaging flow cytometry technology (ImageStreamX system – ISX) (Pugsley, 2017). The quantification of dots of GFP-LC3 per cell shows that OMVs HlyF WT treatment induces a 5-fold increase of LC3-positive structures in eukaryotic cells compared to non-treated cells (**Fig. 2A –B**). We also showed that the accumulation of LC3-positive structures increases substantially with the concentration of OMVs HlyF WT until it reaches a plateau from a concentration of 4μg/mL of OMVs (**Fig. 2C-D**). A dose effect showed that no matter how much OMVs of the non-HlyF producing strain are put on the HeLa cells, no LC3-positive structures accumulation is ever observed (**Fig. 2C-D**). Finally, measurement of the level of LC3-II at different time points in HeLa cells showed that autophagosome increase in HeLa cells lasted at least 72 hours after the beginning of OMV HlyF WT treatment (**Suppl. Fig. 2**).

**Figure 2.**
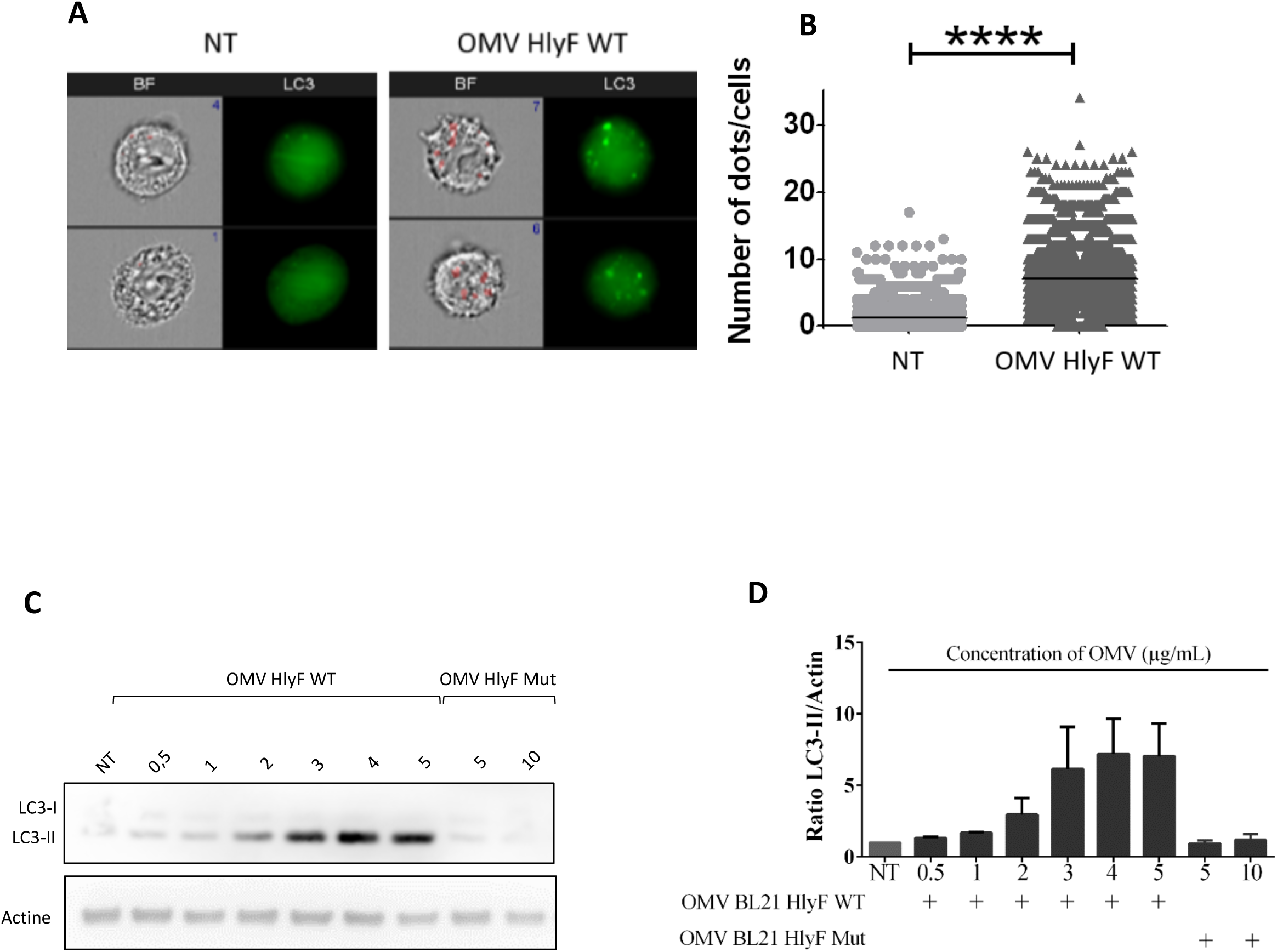
OMVs-induced increase of LC3-positive structures numbers is dose-dependent. (**A**) Images acquired by ISX of GFP-LC3 HeLa cells treated 3hr with OMV from BL21 HlyF WT 5μg/mL. On the left panel, brightfield channel (BF) images the cells according to their size and morphology; on the right panel, LC3 channel images cells according to their fluorescence in GFP-LC3 in green. Number of LC3 green dots of LC3-GFP per cell is indicated in blue in the dial at the top right of the BF channel. The mask is shown in red on the BF. Scale bars = 7μm. Images representatives of 4 independent experiments. (**B**) Quantification of (A). Number of LC3 green dots of LC3-GFP per cell counted with the Spot count function is indicated on the vertical axis on a minimum of 3000 cells per condition. Each square represents a cell image acquired by ISX. The graphs show the mean for each condition. ****p<0.0001 t test. Graph representative of 4 independent experiments. (**C**) Hela cells were treated with increasing concentrations of OMVs from BL21 HlyF WT for 3hr then LC3 and actin were analyzed by western blot. Experiment reproduced 3 times independently. (**D)** Quantification of the LC3B-II/actin ratios of the Fig. 2C obtained by densitometric analysis of 3 independent experiments. The graph shows the mean and the standard deviation for each condition.

### *OMVs from HlyF-expressing* E. coli *triggers the specific accumulation of autophagosomes in host cells*

Accumulation of LC3-positive structures in eukaryotic cells following OMVs treatment could result from an accumulation of autophagosomes or from LC3-associated phagocytosis vesicles (LAP). To distinguish between these two types of vesicles, we first observed that OMVs were not sequestered inside the LC3-positive vesicles, as there is no co-localization between GFP-LC3 structures and OMV stained with DiI (**Fig. 3A**), suggesting that the accumulation of LC3-positive structures does not result from LAP and the phagocytosis of OMVs prior to their degradation but from autophagosomes. We confirmed this result by transmission electron microscopy analysis of Hela cells treated by OMVs, in which we observed the accumulation of double-membrane vesicles, indicative of autophagosome, rather than single-membrane vesicle of LAP (**Fig. 3B**).

**Figure 3.**
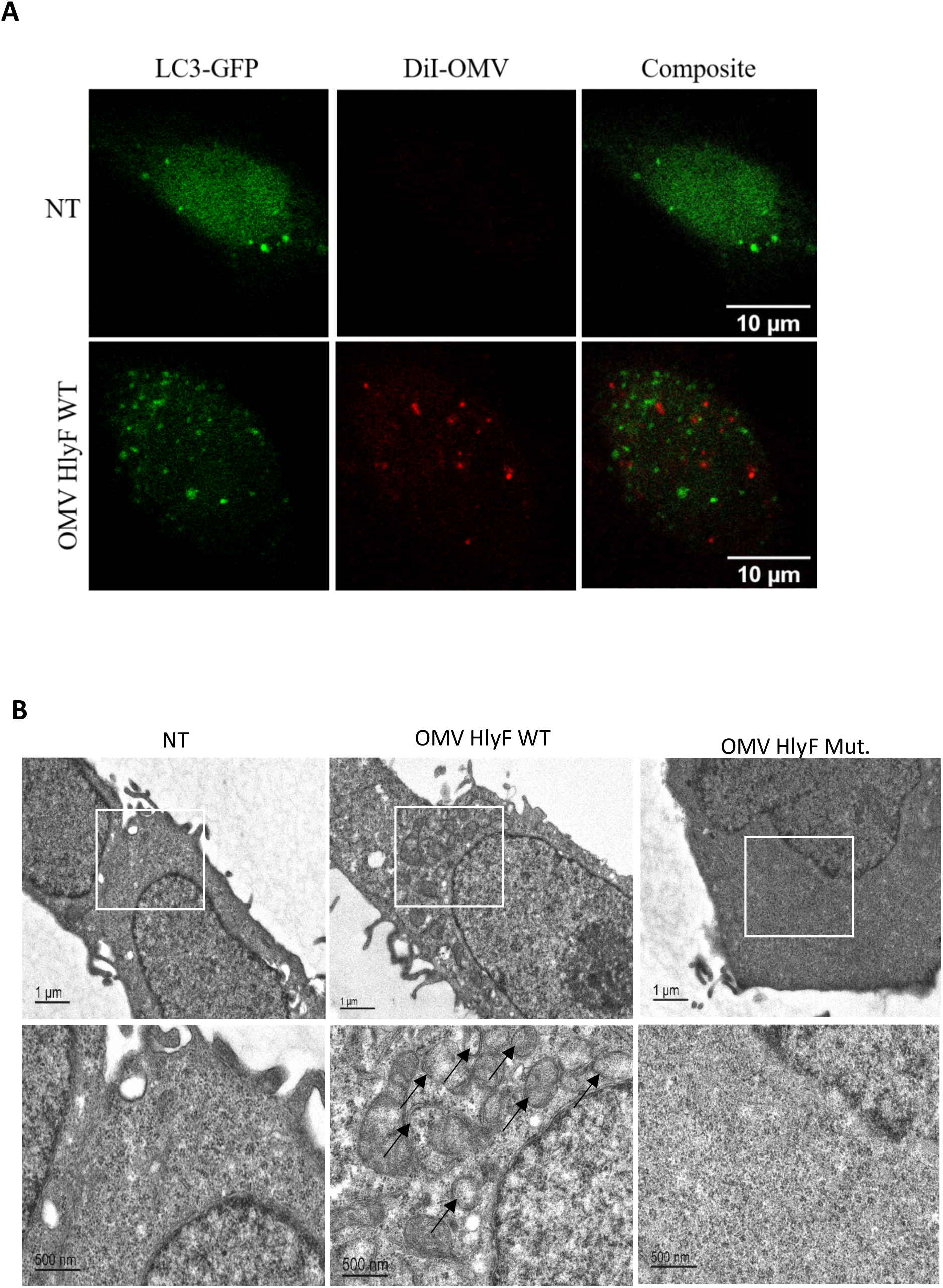
OMVs HlyF WT triggers the specific accumulation of autophagosomes in host cells. (**A**) Confocal images of LC3-GFP (green) and fluorescently labelled OMVs with DiI (Red) fluorescence in LC3-GFP Hela cells. The cells were incubated with OMVs from BL21 HlyF WT 5μg/mL for 3hr. 100 cells were analyzed per condition. Scale bar = 10μm. Images representative of 3 independent experiments. (**B**) Ultrastructure of Hela cells visualized by TEM. The cells were incubated with OMVs from BL21 HlyF WT or from BL21 HlyF Mut 5μg/mL for 3hr. 15 cells were analyzed per condition. Scale bar = 1μm (upper panel) and scale bar = 500nm (bottom panel). Arrows indicate double-membrane autophagosomes. The bottom images are a magnification of the part of the top images within the rectangles.

### *OMVs from HlyF-expressing* E. coli *induce an autophagic blockade*

Autophagy is a dynamic process in which the number of autophagosomes depends on their formation and further degradation. This is referred as autophagic flux. Thus, OMV-dependent accumulation of autophagosomes could result either from an increased synthesis of autophagosomes or from a decreased degradation following their fusion with lysosomes (**Fig. 4A**) (Mizushima et al., 2010; Muñoz-Braceras and Escalante, 2016). To decipher the cellular basis of increase of autophagosomes number after OMV treatment, we monitored the level of autophagosomes-associated LC3-II in HeLa cells. As expected, LC3-II level increased when lysosomal-dependent LC3-II degradation is blocked by the lysosomal inhibitor chloroquine. However, these levels were comparable in cells treated with OMVs in presence of chloroquine (**Fig. 4B, lines 2 and 4, quantified in Fig. 4C**). Absence of cumulative effect of OMVs and chloroquine on LC3-II level suggests that OMVs HlyF WT induce an autophagy blockade in HeLa cells similarly to chloroquine treatment. These results were confirmed by quantification of GFP-LC3 cells treated by OMVs HlyF WT in presence or absence of chloroquine by imaging flow cytometry.

**Figure 4.**
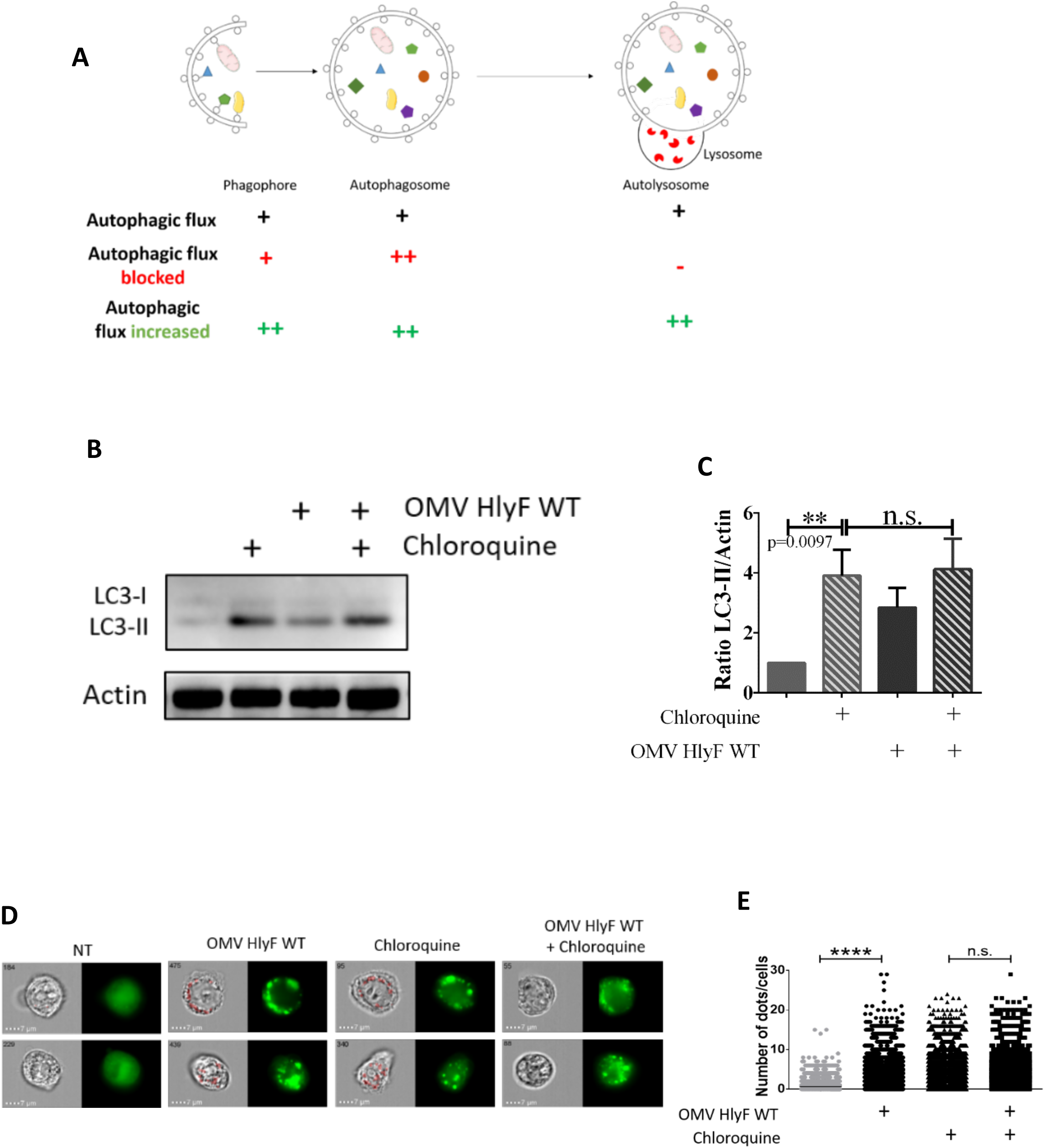
Autophagy flux is blocked by OMVs HlyF WT. (**A**) Schema of the autophagic flux and of the accumulation of different autophagic structures depending on the autophagic dynamic regulation (adapted from Mizushima et al., 2010). (**B**) Hela cells were treated with OMVs from BL21 HlyF WT 5μg/mL and/or 50μM chloroquine for 3hr before LC3 and actin western-blot analysis. Experiment reproduced 3 times independently. (**C**) Quantification of the LC3B-II/actin ratios of the Fig. 4B obtained by densitometric analysis of 3 independent experiments. The graph shows the mean and the standard deviation for each condition. **p = 0.0097 t test, n.s. = non-significant. (**D**) LC3-GFP Hela cells were treated with OMVs from BL21 HlyF WT 5μg/mL and/or 50μM chloroquine for 3hr before ISX analysis. On the left panel, brightfield channel (BF) images; on the right panel, LC3 channel image cells according to their fluorescence in GFP-LC3 in green. Scale bars = 7μm. Images representatives of 3 independent experiments. (**E**) Quantification of (C). Number of LC3 green dots of LC3-GFP per cell counted with the Spot count function is indicated on the vertical axis, on a minimum of 3000 cells per condition. Each square represents a cell image acquired by ISX. The graphs show the mean for each condition. Experiment reproduced 3 times independently, ****p<0.0001 t test.

We observed that the number of GFP-LC3 fluorescent dots in OMV-treated cells is not further increased by chloroquine which would have been the case if the autophagic flux had been stimulated (**Fig. 4D-E**). Altogether, these results showed that OMVs HlyF WT treatment induces an autophagic blockade in eukaryotic cells rather than an induction of the flux.

### *OMVs from HlyF-expressing* E. coli *prevents the autophago-lysosome fusion and autophagosomes clearance*

To characterize which step of the autophagy pathway was inhibited, we used the HeLa-Difluo hLC3 cell line. These cells express a fusion protein RFP::GFP::LC3, in which the N-terminus of LC3 is fused to two fluorescent reporter proteins: an RFP (acid-stable) and a GFP (acid-sensitive). Autophagosomes that contain intact RFP::GFP::LC3 proteins emit both RFP and GFP signals resulting in a co-localization of GFP- and RFP-positive puncta. After fusion of the autophagosomes with the lysosomes, the GFP fluorescence diminishes due to the acidification of the autophagolysosome, while the red fluorescence corresponding to the remaining acidic -resistant RFP-LC3 signal is maintained (Kimura et al., 2007; Loos et al., 2014). We performed confocal imaging of HeLa-Difluo hLC3 cells. As expected, in the control condition treated with lysosomal inhibitor chloroquine, we observed an accumulation of co-localization of GFP- and RFP-positive puncta corresponding to autophagosomes harboring GFP and RFP LC3. When cells were starved with HBSS, known to induce the autophagic flux, a proportion of GFP fluorescence was decreased by the acidic pH environment while acid-resistant red fluorescence persisted, denoting that fusion of autophagosome with lysosome occurred in these cells. In contrast, in cells treated with OMVs HlyF WT, we observed an increased number of co-localization of GFP- and RFP-positive puncta and absence of red-fluorescent-associated autophagolysosome, indicating that the autophagic blockade occurs before the acidification of the autophagosomal compartments (**Fig. 5A**).

**Figure 5.**
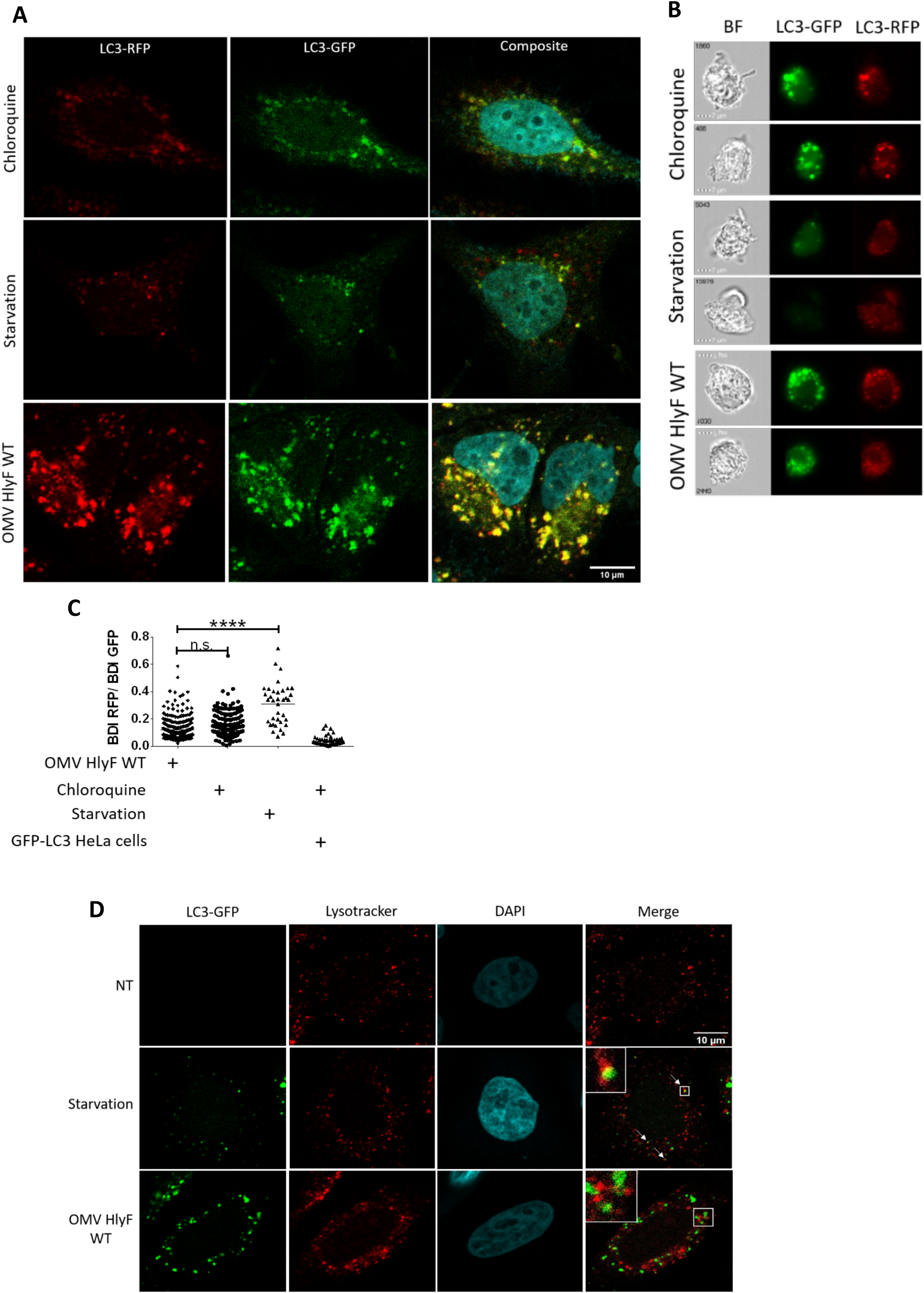
Autophagy flux blocked by OMVs HlyF WT at lysosomal fusion step. (**A**) Confocal images of DiFluo Hela cells deprived with HBSS or treated for 3hr with 50μM chloroquine or with OMVs from BL21 HlyF WT 5μg/mL. Images of LC3-GFP fluorescence (green), LC3-RFP fluorescence (red), and the overlay with DAPI (blue) which shows co-localization of GFP- and RFP-positive puncta. Scale bar = 10μm. 50 cells were analyzed per condition. Images representative of 3 independent experiments. (**B**) Images acquired by ISX of DiFluo Hela cells deprived with HBSS or treated for 3hr with 50μM chloroquine or with OMVs from BL21 HlyF WT 5μg/mL. On the left panel, brightfield channel (BF), on the medium panel, LC3-GFP channel images cells according to their fluorescence in GFP in green and on the right panel LC3-RFP channel images cells according to their fluorescence in RFP in red. Scale bars = 7 μm. Images representatives of 3 independent experiments. (**C**) Quantification of BDI (GFP and RFP) by ISX of DiFluo Hela cells deprived with HBSS or treated for 3hr with chloroquine 50μM or with OMVs from BL21 HlyF WT 5μg/mL. Each square represents the ratio of the intensity of RFP and GFP within a cell. The graph shows the mean of each condition. Experiment reproduced 3 times independently. ****p< 0.0001 t test, n.s. = non-significant. (**D**) Confocal images of LC3-GFP (green), LysoTracker Red (red) and DAPI (blue) fluorescence in LC3-GFP Hela cells. The cells were incubated with OMVs from BL21 HlyF WT 5μg/mL for 3hr and LysoTracker Red for 1hr. 100 cells were analyzed per condition. Scale bar = 10μm. Arrows indicates colocalization foci. Images representative of 3 independent experiments.

To confirm this result, we quantified the level of fluorescently labeled RFP and GFP LC3 in HeLa-Difluo hLC3 by imaging flow cytometry. In contrast to starvation but similarly to chloroquine treatment, we observed that in cells treated by OMVs the ratio RFP/GFP was low, indicating that the GFP is preserved from autophagosome acidification (**Fig 5B-C**).

Finally, we stained acidic organelles in GFP-LC3 cells with LysoTracker Red labeling. We checked that upon starvation, GFP-LC3 Hela cells harbored co-localization of GFP- and RFP-positive puncta (highlighted by white arrows) resulting from the co-localization of autophagosomes and lysosomes merged signals. In contrast, absence of co-localization of GFP- and RFP-positive puncta indicated that OMVs HlyF WT treatment of GFP-LC3 HeLa cells prevented lysosomal acidification of autophagosomes (**Fig. 5D**). These results showed that autophagy flux is blocked by OMV at lysosomal fusion step. Moreover, persistance of signal corresponding LC3-II up to 72hr after OMV addition (suppl. Fig. 2) supports that autophagosomal clearance is inhibited.

### *OMVs from HlyF-expressing E. coli induce a stronger activation of the non-canonical inflammasome pathway compared to OMVs from* E. coli *expressing a non-functional HlyF protein*

Numerous studies showed that autophagy is a negative regulator that prevents excessive activation of inflammasomes (Biasizzo and Kopitar-Jerala, 2020; Deretic et al., 2013; Harris et al., 2017; Krakauer, 2019; Netea-Maier et al., 2015; Qian et al., 2017; Saitoh et al., 2008; Shi et al., 2012). We checked that OMVs HlyF WT cause a substantial autophagosomes accumulation in bone marrow-derived macrophages (BMDM) cells (**Suppl. Fig. 3A-B**) and in the THP1 cell line (**Suppl. Fig. 3C-D**). We therefore speculated that OMV-dependent inhibition of autophagic flux could exacerbate inflammasome activation. OMV-treatment of unprimed BMDM from wild type mice led to cell death (measured by lactase dehydrogenase (LDH) release) and to the excretion of mature IL-1β in the supernatant of cell culture. Importantly, these phenotypes were significantly exacerbated when BMDM were treated with OMVs HlyF WT compared to OMVs produced in bacteria expressing inactive HlyF. These phenotypes were dependent of the activation of the non-canonical inflammasome pathway since they are abolished in inflammasome-deficient BMDM invalidated for caspase 11, caspase 1/11 or gasdermin D (**Fig. 6A**). Furthermore, we deciphered the impact of autophagic blockage on the inflammasome activation following OMV treatment, using 2 different autophagic inhibitors (wortmannin and bafilomycin). As expected, autophagic inhibition alone does not activate the inflammasome machinery. Interestingly, we observed that autophagic inhibition by bafilomycin and wortmannin abolished the autophagic negative feedback loop and thus, restored the overactivation of the inflammasome in OMV HlyF Mut treated cells to the level of OMV HlyF WT treated BMDM (**Fig. 6B-C**). In accordance with the literature, these results suggest that the specific exacerbation of the non-canonical inflammasome pathway activation induced by OMVs HlyF WT treatment was due to their ability to block the autophagic flux (Biasizzo and Kopitar-Jerala, 2020; Deretic et al., 2013; Harris et al., 2017; Krakauer, 2019; Netea-Maier et al., 2015; Qian et al., 2017; Saitoh et al., 2008; Shi et al., 2012). Finally, since activation of the non-canonical inflammasome pathway is related to the sensing of LPS through host guanylate-binding proteins in the cytosol (Santos et al., 2018), we transfected purified LPS from OMVs HlyF WT and from OMVs HlyF Mut directly into the cytosol of BMDM. We observed that the activation of the non-canonical inflammasome pathway, measured by cell death and IL-1β release, was similar regardless of the LPS transfected (**Fig. 6D**). This result suggests that the overactivation of the non-canonical inflammasome pathway does not depend on LPS difference but rather results from OMV HlyF WT ability to block the autophagy.

**Figure 6.**
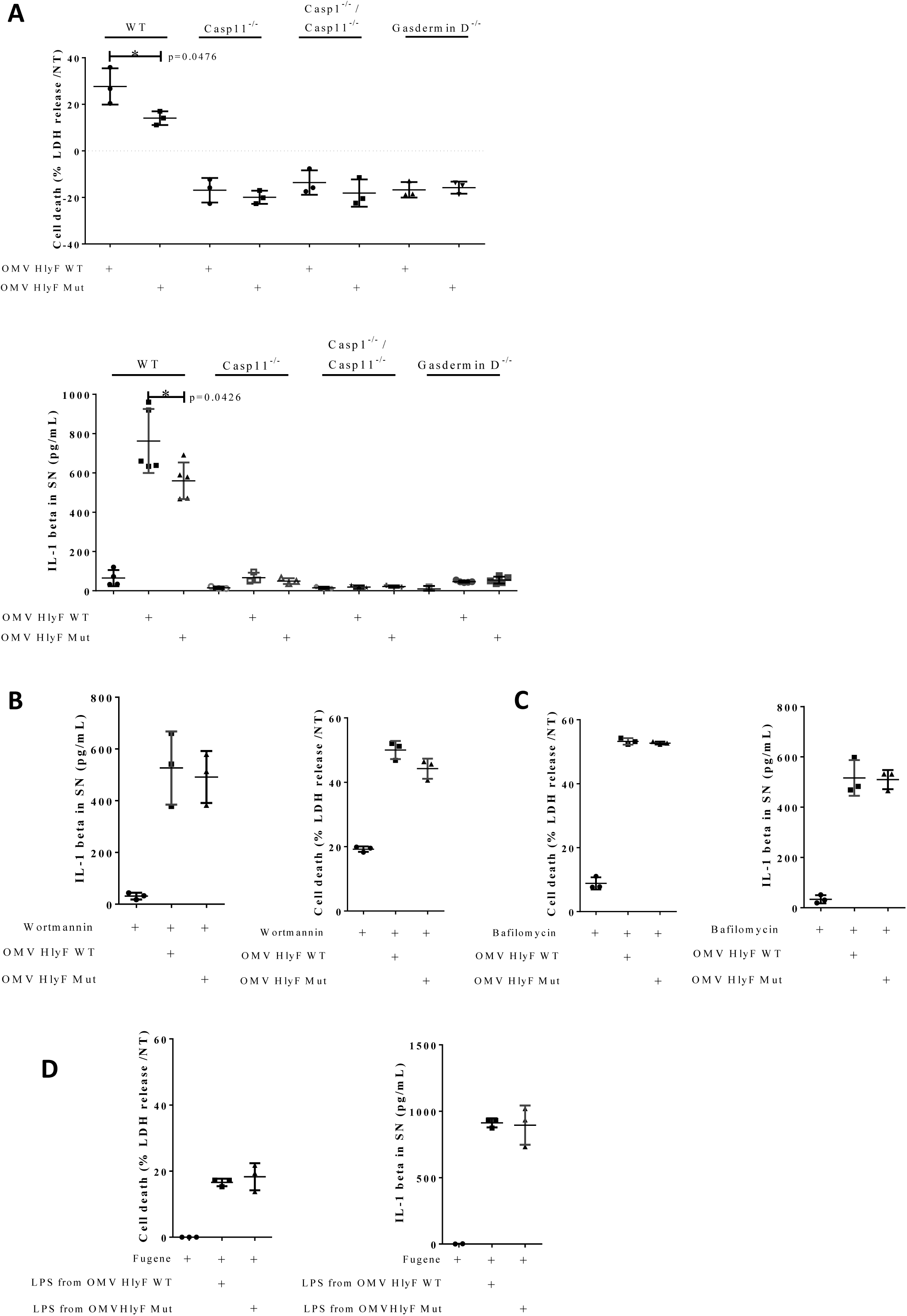
OMVs HlyF WT induce a stronger activation of the non-canonical inflammasome pathway compared to OMVs HlyF Mut. (**A**) Release of LDH and IL-1β from unprimed WT, Casp11^-/-^, Casp1^-/-^ /Casp11^-/-^ and Gasdermin D^-/-^ mice BMDMs treated for 24 hr with OMV HlyF WT or OMV HlyF Mut 5μg/mL. The graphs show the mean and the standard deviation of at least 3 independent experiments for each condition. Each dot represents the value obtained in one experiment. *p<0.05 t test. (**B**) Release of LDH and IL-1β from unprimed WT mice BMDM treated for 24 h with OMV HlyF WT or OMV HlyF Mut 5μg/mL with 10μM wortmaninn. The graphs show the mean and the standard deviation of 3 independent experiments for each condition. Each dot represents the value obtained in one experiment. (**C**) Release of LDH and IL-1β from unprimed WT mice BMDM treated for 24 h with OMV HlyF WT or OMV HlyF Mut 5μg/mL with 10 nM bafilomycin. The graphs show the mean and the standard deviation of 3 independent experiments for each condition. Each dot represents the value obtained in one experiment. **(D**) Release of LDH and IL-1β from unprimed WT mice BMDM transfected with LPS purified from OMVs BL21 HlyF Mut or from OMVs BL21 HlyF WT (1μg/mL using FuGENE HD, Promega) for 24hr. The graphs show the mean and the standard deviation of 3 independent experiments for each condition. Each dot represents the value obtained in one experiment.

## Discussion

In this study, we confirmed that pathogenic E. coli strains harboring a virulence plasmid coding for hlyF overproduce OMVs that modulate autophagy (Murase et al., 2016). However, we showed that this modulation was due to an autophagic blockage instead of an induction of the flux as initially thought (Murase et al., 2016). Moreover, we demonstrated that OMVs HlyF WT block autophagosome maturation and fusion with lysosomes and inhibit their clearance from host cytosol. Autophagy is recognized as an innate antibacterial mechanism by direct lysosomal degradation of intracellular pathogen. After engulfment of the cytosolic pathogen, autophagosomes fuse with lysosomes, leading to degradation of the sequestered content (Netea-Maier et al., 2015). Thus, autophagy that is crucial to maintain cellular homeostasis, acts also as the first line of host defense against bacterial pathogen as it fights against the intruder very early during infection (Losier et al., 2019). It has been shown recently that some ExPEC strains hijack the host autophagy machinery to promote their intracellular survival and replication in macrophages (Zhuge et al., 2018). This ability is due to the inhibition of the fusion of ExPEC-containing phagosomes with lysosomes and their subsequent degradation (Zhuge et al., 2018). This observation is consistent with the properties we demonstrated here, i.e. OMVs preventing the phagolysosomal fusion and acidification in macrophages. In line with this idea, the fact that the adherent-invasive E. coli (AIEC) reference strain NRG 857Cc harbors the hlyF gene suggests that OMVs actively participate in the virulence of these pathogenic bacteria by favoring their survival and their replication into eukaryotic cells (McPhee et al., 2014). Pathogenic bacteria could exploit HlyF OMV property to block their autophagosomal clearance into epithelial cells and macrophages.

Thus, the contribution of OMVs produced by HlyF-expressing E. coli to bacterial virulence includes autophagic blockage but also the dissemination and delivery of their virulence factors potentially far from the source of infection, directly into the cytoplasm of target cells. In line with this idea, OMVs play a very important role in mediating the cytosolic localization of lipopolysaccharide (LPS), an endotoxin that is present at their surface. Once in the host cytosol, LPS is recognized by the non-canonical inflammasome machinery and thereby activates an inflammatory immune response (Kaparakis-Liaskos and Ferrero, 2015; Santos et al., 2018; Vanaja et al., 2016). However, excessive inflammation activation by LPS can be deleterious for the host. The inflammatory burst they cause can also lead to organ dysfunction which can worsen up to sepsis in severe cases (Chen et al., 2018; Schroder et al., 2018). At the early stage of sepsis, macrophages release large amounts of pro-inflammatory molecules that exacerbate inflammatory response. It is then followed by an immunosuppression due to the massive apoptosis of macrophages, which makes host susceptible to recurrent infections (Cecconi et al., 2018; van der Poll et al., 2017). The role of OMVs in sepsis is highlighted by the fact that injection of OMVs alone in mice recapitulates sepsis-associated exacerbated inflammation and mortality (Eren et al., 2020). Thus, this inflammation machinery initiated by OMV recognition must be tightly regulated. Here, we show that OMV HlyF WT provoke an exacerbated activation of the non-canonical inflammasome activation and IL-1β release, one of the main mechanisms through which innate and adaptive immune responses are induced by an infection (Eren et al., 2020; Gomes and Dikic, 2014; Schroder et al., 2018). We showed that not only OMVs HlyF WT induces inflammation actions, they also block autophagy, a major negative regular of excessive inflammation activation and IL-1β release (Crişan et al., 2011; Saitoh et al., 2008). In particular, autophagy induction prevents the onset of sepsis (Biasizzo and Kopitar-Jerala, 2020). This negative regulation mainly relies through effects on the secretion of immune mediators and on the removal of endogenous inflammasome agonists such as PAMPs (Deretic et al., 2013; Harris et al., 2017; Krakauer, 2019; Netea-Maier et al., 2015; Qian et al., 2017). Therefore, not only OMVs favor intracellular delivery of LPS and PAMPs into cells but inhibition of autosomal clearance of these inflammasome activators by OMV HlyF WT impairs inflammasome negative feedback control, leading to an excessive and uncontrolled inflammatory condition detrimental for the host.

Our study unravels the mechanisms by which HlyF expression by E. coli confers hyper-virulent properties to OMVs, as they gain the property to manipulate the host cell autophagic and inflammatory responses to bacterial benefit both at the site and at great distance from the site of infection. An additional contribution of OMVs HlyF to bacterial virulence could be associated to the dissemination of toxins (Bielaszewska et al., 2017; Murase et al., 2016). Overall, OMVs HlyF contribute significantly to the virulence of pathogenic bacteria.

## Materials and methods

### Bacterial strains and growth conditions

SP15 is an extra-intestinal pathogenic *E. coli* (ExPEC) strain isolated from a neonatal meningitis case (Johnson et al., 2002). SP15, SP15Δhlyf and SP15ΔhlyfGST-hlyF were previously described in (Murase et al., 2016). pGEX6P-1 plasmids expressing GST-HlyF protein wild type or inactivated by a mutation on its putative catalytic site (**Suppl. Fig. 1A**) were transformed *in E. coli* BL21(DE3) strain. SP15 and BL21(DE3) strains and derivatives were inoculated and grown in Terrific Broth with appropriate antibiotics for 8hr at 37°C with agitation.

### Purification of bacterial OMVs

After 8hr of culture the bacterial suspension was centrifuged at 6500g for 10min at 4°C. The supernatant was filtered (pore size 0.45μm) to obtain bacterial-free supernatant. The filtered supernatant was ultrafiltered and concentrated using tangential concentrator and PES membrane cassette (pore size, 100kDa, Sartorius) and ultracentrifuged at 150,000g for 1.5hr at 4°C. After removing the supernatant, pelleted OMV were resuspended in sterile PBS with Ca^2+^ and Mg^2+^ and stored at 4°C. To visualize the concentration of OMVs and the purity of the preparation, negative staining transmission electron microscopy (TEM) was performed according to standard procedures. The concentration of OMVs in the suspension correlates with protein concentration evaluated by BCA protein assay (DC™ protein assay Biorad) (**Suppl. Fig. 1C**). LPS concentration in OMV was evaluated by Purpald assay as described in (Lee and Tsai, 1999). **Values were normalized to glycerol concentration. OMV diameter was calculated by dynamic light scattering using Zetasizer Nano instrument (Malvern) (Suppl. Fig. 1B)**.

OMVs in PBS were fluorescently labeled using 1% (v/v) DiI (Thermofisher) for 30 min at 37°C.

### Purification and transfection of LPS from OMVs

LPS was extracted from purified OMV using LPS extraction kit (iNtRON Biotechnology) according to manufacturer’s instruction. LPS concentration was determined by Purpald assay as described in (Lee and Tsai, 1999). Transfection of cells with purified LPS was done at a concentration of 500ng/50,000 cells, using FuGeneHD (Promega) transfection reagent in Opti-MEM, as previously described (Santos et al., 2018).

### Eukaryotic cell culture and treatments

HeLa cells, HeLa-LC3-GFP cells and *HeLa-Difluo* hLC3 cells (Invivogen) were cultured in DMEM with Glutamax (Sigma) supplemented with 10% fetal bovine serum (FBS). *HeLa-Difluo* hLC3 cells were maintained in growth medium supplemented with the selection antibiotic *Zeocin 200μg/mL (Invivogen)*. One day before the experiment, HeLa cells were seeded at a density of 5 × 10^4^ cells per mL in DMEM supplemented with 10% FBS. OMV treatment were done by adding directly in the medium of cell culture the volume of OMV corresponding to the amount of proteins indicated. The cells were collected 3h after OMV addition.

THP1 cells were cultured in RPMI with Glutamax (Sigma) supplemented with 10% FBS. One day before the experiment, THP1 cells were seeded at a density of 3 × 10^5^ cells per mL. OMV treatment were done by adding directly in the medium of cell culture the volume of OMV corresponding to the amount of proteins indicated. The cells were collected 3h after OMV addition.

Bone marrow-derived macrophages (BMDM) were prepared from fresh bone marrow isolated from WT, Caspase 11^-/-^, Caspases 1^-/-^/11^-/-^ or Gasdermin D^-/-^ deficient mice. Bone marrow cells were allowed to differentiate in macrophages by incubation for 7 days in DMEM supplemented with 10% fetal calf serum (Bioconcept), 25ng/ml M-CSF (Immunotools), 10mM HEPES, and nonessential amino acids (Life Technology). One day before treatment, BMDM were seeded into 24 well plates at a density of 2.5 × 10^5^ cells per well. OMV treatment, were done by adding the volume of OMV corresponding to the amount of proteins indicated in Opti-MEM medium. The cells and supernatant were collected 24hr after OMV addition.

### Immunoblotting

Cells were treated with the OMV or left untreated for the indicated times. Then cells were washed twice with cold PBS, lysed in NuPAGE LDS sample buffer with reducing agent (Invitrogen) and sonicated. Cell extracts were boiled for 5min at 95°C, separated by SDS-PAGE and transferred to PVDF membranes (Invitrogen). The membranes were blocked in 5% BSA in 0.1% Tween-20 in TBS and probed with primary antibodies. The primary antibodies used were Rabbit anti-LC3 (Cell Signaling S 3868), Mouse anti-actin (Abcam Ab 3280). After incubation with Goat anti-Rabbit and Goat anti-Mouse HRP-conjugated secondary antibodies (Promega), proteins were visualized using chemiluminescent peroxidase substrat detection reagents (Sigma), and acquired by the ChemiDoc XRS+ System (BioRad). Densitometric analyses of immunoblots were performed using the ImageJ software.

### Fluorescence microscopy

GFP-LC3 HeLa cells or *HeLa-Difluo* hLC3 cells were seeded onto Nunc Lab-Tek Chamber Slide system 8 wells (Sigma). The day of the experiment, cells were incubated with the indicated treatment or left untreated for the indicated times. LysoTracker Red (Invitrogen) staining was achieved by adding the Lysotracker probe at a final concentration of 50nM directly in the medium of cell culture for 1hr. Then the cells were washed and fixed with 4% paraformaldehyde for 15min on ice. DNA was visualized using Fluoroshield with 4,6-diamidino-2-phenylindole (DAPI, Sigma). Images were acquired using a Zeiss LSM 710 confocal microscope and subsequently processed using the ImageJ software package.

### Transmission electron microscopy

Negative staining TEM for OMV was performed according to standard procedures. Briefly, 5μL of OMV samples were added to carbon coated copper mesh grid and stained with 1% uranyl acetate for 1 minute and imaged with Jeol JEM-1400 transmission electon microscope. For Hela cells ultrastructure visualization, cells were cultured on 6-well plates at a density of 1,5 × 10^6^ cells/well one day prior to the experiment. After OMV treatment, cells were fixed with final concentrations of 2.5% glutaraldehyde / 2% paraformaldehyde in 0.1M Sorensen buffer (pH 7.2) volume/volume in the culture medium for 15 minutes at room temperature. After this primary fixation, the cells were secondarily fixed with 2.5% glutaraldehyde / 2% paraformaldehyde in 0.1M Sorensen buffer (pH 7.2) for 1.15 hr at room temperature. Then cells were rinsed three times in Sorensen buffer for 10 min each time and let in Sorensen buffer with 1% paraformaldehyde at 4°C until they were then scraped off, pelleted, concentrated in agarose, and treated for 1 h with 2% aqueous uranyl acetate. The samples were then dehydrated in a graded ethanol series and embedded in Epon. After 48 h of polymerization at 60 °C, ultrathin sections (80 nm thick) were mounted on 200 mesh Formvar-carbon-coated copper grids. Finally, sections were stained with Uranyless and lead citrate. Grids were examined with a TEM (Jeol JEM-1400, JEOL Inc, Peabody, MA, USA) at 80 kV. Images were acquired using a digital camera (Gatan Orius, Gatan Inc, Pleasanton, CA, USA) and subsequently processed using the ImageJ software package.

### Measurement of LDH release

Cell death was quantified by measuring LDH release, using the CyQUANT™ LDH Cytotoxicity Assay (Invitrogen). To normalize for spontaneous cell lysis, the percentage of cell death was calculated as follows: [(LDH sample) – (LDH non-treated control)] / [(LDH positive control) – (LDH non-treated control)] × 100.

### Measurement of IL-1β release

Quantification of IL-1β secretion in cell supernatant was measured by ELISA (Invitrogen, IL-1 beta Mouse Uncoated ELISA Kit, **#** 88-7013-88), according to the manufacturer’s instructions.

### Imaging Flow Cytometry

GFP-LC3 HeLa cells or HeLa Difluo hLC3 cells were incubated with the indicated treatment or left untreated for the indicated times. Then the cells were washed, trypsinized, fixed with 4% paraformaldehyde for 15min on ice. Image of each cell was acquired while they were run on the **ImageStreamX Mark II (Amnis-EMD Millipore)** at 5 × 10^7^ cells per mL in PBS. In each experiment 5,000 (GFP+) events per sample were acquired (ensuring a sufficient number of events remaining for statistically robust analysis) at 60X magnification. Single stain samples were also collected using the same settings and used as compensation controls to generate compensation matrix.

Image analysis was completed using image-based algorithms in the ImageStream Data Exploration and Analysis Software (IDEAS_6.2.64.0, EMD Millipore). The number of GFP-LC3 puncta per cell was assessed with the feature “Spot Counting”. A mask based on GFP-LC3 staining was employed (SpotCount (M02, LC3, Bright, 16, 2, 1)) to visualize the GFP-LC3 puncta. A feature in the IDEAS software called Bright Detail Intensity R7 (BDI) was used to assess the intensity of bright spots in the cell image that have radii smaller than a threshold size (7 pixels in this study), while neglecting background staining. The ratio of BDI RFP/ BDI GFP brightness was thus assessed. The same mask and feature were automatically applied for each condition.

## Abbreviations

AIEC: adherent-invasive *E. coli*
BDI: Bright Detail Intensity
BMDM: bone marrow-derived macrophages
Casp: Caspase
*E. coli*: *Escherichia coli*
EHEC: enterohemorrhagic *E. coli*
ExPEC: extra-intestinal pathogenic *E. coli*
GSDM-D: Gasdermin-D
GFP: Green Fluorescent Protein
HBSS: Hanks’balanced salt solution
HlyF: Hemolysin F
IL-1β: interleukin-1β
ISX: ImageStreamX system
LPS: Lipopolysaccharide
OMV: outer membrane vesicle
RFP: Red Fluorescent Protein
TEM: transmission electron microscopy
WT: Wild type
Mut: Mutated

## Acknowledgments

We gratefully acknowledge the imaging platform of INFINITy institute. This work also benefited from the assistance of Stephanie Balor and Vanessa Soldan from the Multiscale Electron Imaging platform (METi) of the Centre de Biologie Intégrative (CBI). We also thank Emmanuel Ravet (Invivogen, Toulouse) for providing us *HeLa-Difluo* hLC3 cells, Marie-Charline Blatche (LAAS institute, Toulouse) for assistance in DLS analysis and Miriam Pinilla (IPBS institute, Toulouse) for her valuable help in the preparation of BMDM cells and the LPS transfection experiments. Finally, we are indebted to Dr MS, Dr J-PM and Dr J-PN for their insightful comments on the manuscript.

## Supplementary materials

Suppl. Figure S1

Suppl. Figure S2

Suppl. Figure S3

