## supplemental figures for "Outer membrane vesicles produced by pathogenic strains of *Escherichia coli* block autophagic flux and exacerbate inflammasome activation"

Suppl. Figure S1

**A**

Sequence alignment:

```
Seq type of catalytic domain of SDR family :     *Y***K*
Seq of wild type HlyF                      : 162 EYTRSKA 168
Seq of mutated HlyF                        : 162 EFTRSAA 168
```

**B**

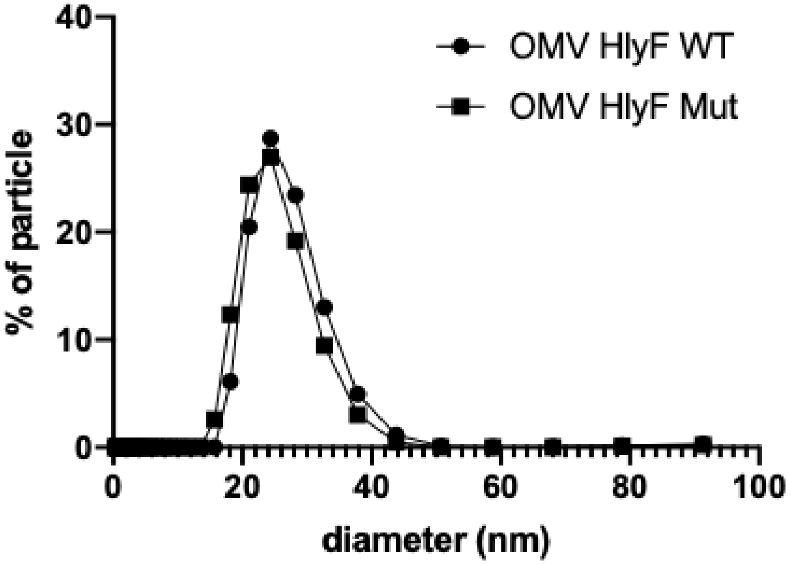


**C**

| | OMV HlyF WT | OMV HlyF Mut |
|---|---|---|
| Diameter, SD (nm) | 25.9, 4.9 | 24.5, 4.8 |
| Protein concentration (mg/mL) | 1 | 1 |
| LPS concentration (mM) | 1.1 | 0.9 |
| Particles concentration (OMV/mL) | $3.10^{11}$ | $3.10^{11}$ |

Suppl. Fig S1: (**A**) Alignment of the catalytic site of short chain reductase (SDR) family (first line), wild type HlyF (second line) and mutated HlyF (third line). Strictly conserved residues (Y and K) and corresponding mutated residues (F and A) are indicated in bold. Amino-acid positions of HlyF are indicated at left and right sides of the sequence. * = any amino-acid. (**B**) Representative curves (from 3 independent experiments) of OMVs BL21 HlyF WT and OMVs BL21 HlyF Mut diameter (expressed as percent of total number) that was calculated by dynamic light scattering (DLS) using Zetasizer Nano (Malvern). (**C**) Characteristics of purified OMVs BL21 HlyF WT and OMVs BL21 HlyF Mut suspension: OMV diameter calculated from panel B, protein concentration (BCA assay), LPS concentration (glycerol equivalent, Purpald assay) and particles concentration are indicated.

Suppl. Figure S2:

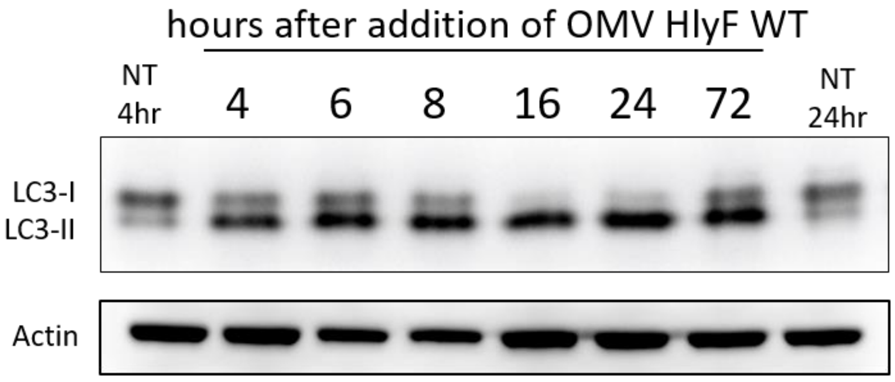


Suppl. Fig. S2: Hela cells were treated with OMV from BL21 HlyF WT 5µg/mL for the indicated time period before LC3 and actin western-blot analysis.

Suppl. Figure S3**:**

**A**

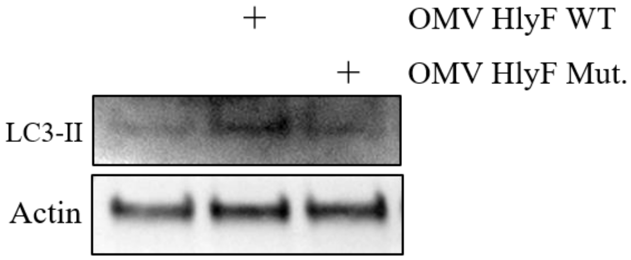


**B**

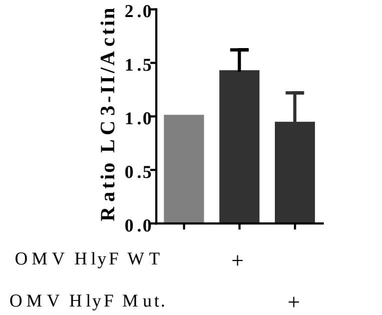


**C**

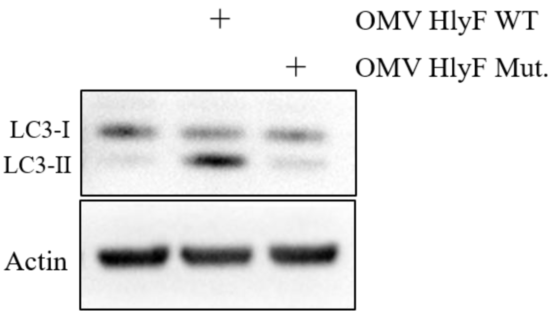


**D**

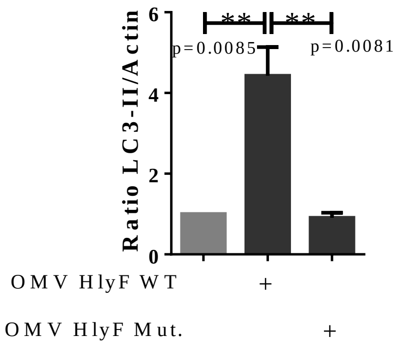


Suppl. Fig S3: (**A**) BMDM from wild type mice were treated with 5µg/mL OMVs from BL21 HlyF WT or from BL21 HlyF Mut for 24hr before LC3 and actin western-blot analysis. Representative of 3 independent experiments. (**B**) Quantification of the LC3B-II/actin ratios of (**A**) obtained by densitometric analysis of 3 independent experiments. The graph shows the mean and the standard deviation for each condition. (**C**) THP1 cells were treated with 10µg/mL OMVs from BL21 HlyF WT or from BL21 HlyF Mut for 24hr before LC3 and actin western-blot analysis. Representative of 3 independent experiments. (**D**) Quantification of the LC3B-II/actin ratios of (**C**) obtained by densitometric analysis of 3 independent experiments. The graph shows the mean and the standard deviation for each condition. **p < 0,01 t test.
